# PCSK9 inhibition protects mice from food allergy

**DOI:** 10.1101/2023.10.10.561796

**Authors:** Victoria Lorant, Martin Klein, Damien Garçon, Thibaud Sotin, Samuel Frey, Marie-Aude Cheminant, Audrey Ayer, Mikaël Croyal, Laurent Flet, Luc Colas, Bertrand Cariou, Grégory Bouchaud, Cédric Le May

## Abstract

The Proprotein Convertase Subtilisin Kexin of type 9 (PCSK9) has been identified in 2003 as the third gene involved in familial hypercholesterolemia. PCSK9 binds to the membrane low-density lipoprotein receptor (LDLR) and promotes its cellular internalization and lysosomal degradation. Beyond this canonical role, PCSK9 was recently described to be involved in several immune responses. However, to date, the contribution of PCSK9 in food allergy remains unknown. Here, we showed that *Pcsk9* deficiency or pharmacological inhibition of circulating PCSK9 with a specific monoclonal antibody (m-Ab) protected mice against symptoms of gliadin-induced-food allergy, such as increased intestinal transit time and ear oedema. Furthermore, specific PCSK9 inhibition during the elicitation steps of allergic process was sufficient to ensure anti-allergic effects in mice. Interestingly, the protective effect of PCSK9 inhibition against food allergy symptoms was independent of the LDLR as PCSK9 inhibitors remained effective in *Ldlr* deficient mice. *In vitro*, we showed that recombinant gain of function PCSK9 (PCSK9 D374Y) increased the percentage of mature bone marrow derived dendritic cells (BMDCs), promoted naïve T cell proliferation and potentiated the gliadin induced basophils degranulation. Altogether, our data demonstrate that PCSK9 inhibition is protective against gliadin induced food allergy in a LDLR-independent manner.

## Introduction

Prevalence of food allergies (FA) significantly increased worldwide over the last thirty years. Currently, 5% to 8% of children and 1% to 2% of adults are concerned by this disease^1,2^. In patients, symptoms may include abdominal pain, vomiting, diarrhea, itchy skin, and/or vascular collapsus^3^. Allergic diseases are classified as the 4^th^ pathology in terms of morbidity by the World Health Organization ^4^ since they impact patients’ quality of life and represent a significant economic burden^5^. However, current therapeutic options remain limited. Therefore, a better understanding of the key immunological mechanisms involved in this process is required to develop new therapeutic strategies. FA is characterized by sensitization mechanism and symptoms elicitation occurrence. Th2 lymphocytes play a key role in sensitization leading to allergen specific immunoglobulin (Ig) E production by B cells^6^. Cross-linking of the allergen-specific-IgE and the Fcε receptors is necessary to activate mast cells/basophils and release pro-inflammatory type 2 molecules such as histamine, interleukin IL-4, IL-5, IL-13 and arachidonic acid derivatives^7^. Recently, it was suggested that blood lipids levels are associated with allergic pathology. In a non-obese pediatric population with atopic predisposition, a positive correlation was found between serum IgE and cholesterol plasma levels^8^. Serum IL-10 concentration is lower in children with dyslipidemia and atopic predisposition than in children with dyslipidemia only^8^. In addition, lipid metabolism was associated with IL-10 synthesis downregulation^9^. In opposite, sterol metabolism and generation of cholesterol precursors are able to promote Th17 cells polarization and IL-6, IL-1, IL-4 and IL-17 release^10^.

Proprotein convertase subtilisin/kexin type 9 (PCSK9) has been identified in 2003 as the third gene involved in familial hypercholesterolemia^11^, a genetic disease associated with high plasma low-density lipoprotein cholesterol (LDL-c) level and premature cardiovascular diseases^12^. PCSK9 is expressed in many organs^13^ but the liver appears to be the sole organ contributing to circulating PCSK9 levels^14^. PCSK9 is a central regulator of cholesterol homeostasis by binding to the extracellular domain of the LDL receptor (LDLR) and thereby promoting its intracellular internalization and lysosomal degradation^15^. In the atherosclerotic plaque, PCSK9 promotes the recruitment of immune cells and confers a pro-inflammatory profile to macrophages, monocytes and smooth muscle cells^16–19^. Other studies determined the effect of PCSK9 on Th17 cells and dendritic cells (DC) maturation and activation in atherosclerotic plaque^20,21^.

Beyond its canonical role, numerous studies reported that PCSK9 could participate in several inflammatory disorders. PCSK9 full deficiency and inhibition confer protection against septic shock response and cytokine cascade^21–25^. PCSK9 inhibition showed protective effects in DSS-induced-colitis rat model^26^. More importantly, it was recently demonstrated that PCSK9 depletion attenuates tumor growth, improves mice survival and increases immune effector cells intra-tumoral infiltration^27^.

Considering the involvement of PCSK9 on immune polarization and the beneficial effect of its inhibition on several inflammatory diseases, we aim to explore the functional impact of PCSK9 deficiency or inhibition on an establish mouse model of wheat-gliadin induced FA.

## Materials and methods

### Animals models

C57Bl6/J, BALB/c (from Charles River laboratories), *Pcsk9^-/-^* and intestinal-specific *Pcsk9* deficient (i*-Pcsk9^-/-^*) ^28^ and *Ldlr*-deficient (*Ldl-r^-/-^*) mice were housed in a 12-hour day/night cycle and had unlimited access to water and gluten-free food. The protocol was approved by the Ethics Committee on Animal Experimentation of Pays de Loire (CEEA-23421).

### Allergy induction protocol

Five-week-old mice were sensitized at day 0, 10 and 20 by intraperitoneal injection of 10 µg deamidated gliadins with aluminium hydroxide (DG/Alum)^29^. At day 27 and day 34, mice were orally challenged with 20 mg of DG in water to induce the allergic reaction. Negative control mice were sensitized with PBS and challenged with water.

### Pharmacological inhibition of PCSK9

Monoclonal anti-PCSK9 antibody (m-Ab, Alirocumab, Sanofi-Aventis, France) was diluted in saline solution and subcutaneously administrated to 4-week-old BALB/c mice (50 mg/kg BW every ten days).

### Intestinal transit time measurement

Intestinal transit time was evaluated at day 27 by measuring the time period between an oral gavage with a solution containing DG (20mg) and red carmin (150 mg/mL) and the appearance of the first red fecal pellet.

### Measurement and histological visualization of ear thickness.

During second challenge, ears thickness was determined with a digital micrometer (Mitutoyo, America Corporation, Aurora, IL) before and one hour 20mg DG gavage^30^. Ears were then harvested, fixed in 4% paraformaldehyde, embedded in paraffin, cut and stained with hematoxylin eosin. 5 µm sections were scanned using the NanoZoomer slide scanner (Hamamatsu Photonics, Japan) and the dermis thickness was quantified using NDP view 2 software (Hamamatsu Photonics K.K., Shizuoka, Japan).

### Isolation of cells and flow cytometry

Mesenteric lymph nodes (MLNs) and spleen were collected and homogenized in in RPMI-1640 medium (Gibco, Thermo Fisher Scientific). 1.10^6^ cells were transferred to a 96-well round-bottom plate and stimulated for 5 h with 50 ng/mL of phorbol-12-myristate-13-acetate and 1 µg/mL of ionomycin (Sigma-Aldrich, Saint Quentin Fallavier, France) with brefeldin A (10 µg/mL, BD Biosciences, Le Pont de Claix, France). Cells were stained with surface markers (CD3-APCH7, CD4-PEPCy7, CD8-PerCP5.5, CD25-BV510, CD9-APC, CD19-PerCP5.5, IgE-BV510, IgD-PE) in the presence of Fc blocker mouse CD16/32 antibody (BD Biosciences). For intracellular staining, the cells were fixed and permeabilized using a Cytofix/Cytoperm kit (BD Biosciences, Le Pont de Claix, France) and stained with FoxP3-APC, IL10-PE, RORγT-BV421, Gata3-BV421, INFγ-PE and IL4-APC. Cells were analyzed on a Fortessa cytometer (BD Biosciences, Le Pont de Claix, France). Data were acquired using Diva software and analyzed with FlowJo v10 (FlowJo LLC, Ashland, OR, USA).

### *In vitro* measurement of PCSK9 effect on dentritic cell maturation, T cell proliferation and basophil degranulation

Detailed methods are provided as supplemental document.

### Statistical analysis

Data were analyzed using GraphPad Prism (La Jolla, CA, USA). Normality and equality of variance were checked by normality test and F-test, respectively. Values were expressed as the mean ± standard error of the mean (SEM) and compared using Mann-Whitney *U* test, one-way ANOVA with Kruskal-Wallis multiple comparison test or two-way ANOVA multiple comparison test. A *p*-value of less than 0.05 was considered as statistically significant.

## Results

### Total, but not intestine-specific, Pcsk9 deficiency protects mice against deamidated gliadin-induced FA symptoms and inflammation

We first explored the impact of full PCSK9 deficiency on FA symptoms (**Fig.1A**). As expected, *Pcsk9^-/-^* mice had a reduced plasma cholesterol level (0.86 ± 0.08 g/L vs 1.20 ± 0.15 in *Pcsk9^+/+^*mice, *P*<0.05) (**Fig.1B**) and an absence of circulating PCSK9 (**Fig.1C**). Previous studies showed that inflammation and allergy can alter plasma PCSK9 and cholesterol levels respectively^31,32^. In our set-up, we observed that plasma cholesterol levels were not affected by FA in *Pcsk9^+/+^* mice (**Fig.1B**), despite a significant rise in plasma PCSK9 level (**Fig.1C**). Hypocholesterolemia was less marked in *Pcsk9^-/-^* mice under FA conditions (**Fig.1B**). We compared the development of FA symptoms in *Pcsk9^+/+^ and Pcsk9^-/-^* mice (**Fig.1A**). Allergic *Pcsk9^+/+^* mice display an increased transit time compared to control *Pcsk9^+/+^* mice (156 ± 11 vs 270 ± 14 min, *P*<0.0001) (**Fig.1D**). No difference was detected between control and allergic *Pcsk9^-/-^* mice (**Fig.1D**). Consistently, we also observed a significant increase of ear thickness in allergic *Pcsk9^+/+^* mice (**Fig.1E**), while allergic *Pcsk9^-/-^* mice did not develop ear oedema (**Fig.1E**). Together, these data indicate that full *Pcsk9* deficiency protects mice against food allergic symptoms development. To determine the cellular mechanisms involved in the anti-allergic effect of PCSK9 deficiency, we measured immune cell populations from MLNs and spleen by flow cytometry. While MLN Th2 and Th17 lymphocytes number were increased by FA in *Pcsk9^+/+^*mice, we did not observe a similar induction in allergic *Pcsk9^-/-^*mice (**Fig.1F and G).** Th2 and Th17 cells from spleen are not modified by FA or PCSK9 deficiency (**S1A ans B**). IgE^+^ B cells number from MLNs were not increased with FA (**Fig.1H**). By contrast, splenic IgE^+^ B cells number was significantly increased in allergic *Pcsk9^+/+^* but not in allergic *Pcsk9^-/-^*mice (**Fig.1I**). Finally, we observed a non-significant trend for a plasma IgG1 increase in *Pcsk9^+/+^* FA mice (**Fig.1J**). Altogether, these results confirm that *Pcsk9* deficiency attenuates the allergic phenotype of FA mice.

**Figure 1.**
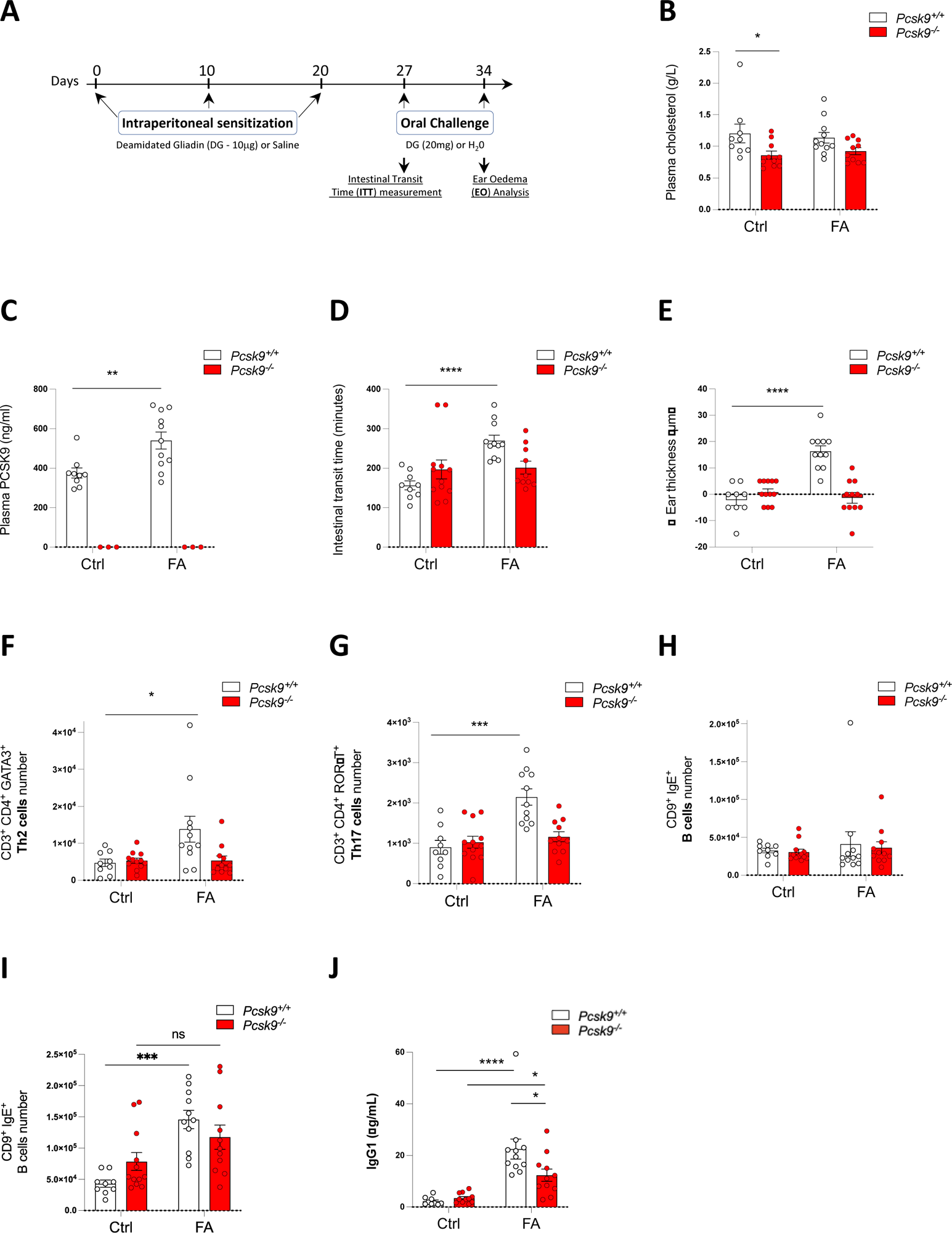
Total PCSK9 deficiency protects mice against deamidated gliadin-induced food allergy. (A) Schematic representation of the food allergy protocol: Five weeks old *Pcsk9^+/+^* and *Pcsk9^-/-^* mice were sensitized at day 0, 10 & 20 with either PBS (Ctrl group) or deamidated gliadins (DG: 50 µg in 200 µL (PBS + alum); Food allergy-FA-Group). One week after the last sensitization, mice were orally challenged with either water (Ctrl) or DG (20 mg in 200 µL of water; FA group). (B) Intestinal transit time measured at day 27 after the first challenge. (C) At day 34, after the second challenge, we assessed the gliadin induced ear oedema. We measured (D) the plasma cholesterol and (E) the plasma PCSK9 levels. Th2 (F) and Th17 lymphocytes (G) and IgE^+^ B cells (H) numbers from mesenteric lymph nodes were quantified by flow cytometry. (I) The number of splenic IgE^+^ B cells was measured by flow cytometry from *Pcsk9^+/+^* and *Pcsk9^-/-^* mice. (J) Plasma IgG1 concentrations were measured by ELISA from plasma harvested 1h after the second elicitation. Data are represented as the mean ± SEM. P-value was determined using Mann-Whitney *U* test; \**P* < 0.05, \*\**P* < 0.01, \*\*\**P* < 0.001 and \*\*\*\**P* < 0.0001. n=9-12 mice per group.

As PCSK9 is highly expressed in the small intestine^14,33^, we verified whether intestinal PCSK9 could be involved in the regulation of FA. The same experimental protocol was reproduced in intestine-specific PCSK9 knock-out (i*-Pcsk9^-/-^) ^-^*mice^28^ (**Fig.S2A)**. Both the intestinal transit time and the ear oedema were similarly increased in i*-Pcsk9^+/+^* and i-*Pcsk9^-/-^* mice following FA (**Fig.S2B-E).** Altogether, these data demonstrate that full *Pcsk9* deficiency protective action does not involve intestinal PCSK9.

### Pharmacological inhibition of circulating PCSK9 by m-Ab reduces food allergy symptoms

To determine the importance of circulating PCSK9 on FA regulation, we blocked its activity using a human PCSK9 m-Ab (**Fig.2A**). Due to the blockade of PCSK9 activity toward LDLR, plasma cholesterol levels were decreased by 40 % following PCSK9 m-Ab treatment (**Fig.2B**). Plasma total PCSK9 levels were increased due to PCSK9 m-Ab induced plasma sequestration (**Fig.2C**). Upon allergic conditions, vehicle-treated BALB/c mice demonstrated a significantly increased intestinal transit time (**Fig.2D**) and increased ear thickness (**Fig.2E**) compared to control mice. The treatment with PCSK9 m-Ab prevented the development of allergic symptoms (**Fig.2D and E**). We measured a significant rise of Th2 and Th17 cells in MLNs of vehicle-treated allergic mice but not in PCSK9 m-Ab treated mice (**Fig.2F and G**). In MLNs, FA strongly increased strongly IgE^+^ cells number in vehicle treated mice compared to non-allergic mice (**Fig. 2H**). Under PCSK9 m-Ab treatment, allergy did not induce a significant increase of the MLNs IgE^+^ cells number. In the spleen, we did observe similar patterns (**Fig.S3A-C**).

**Figure 2.**
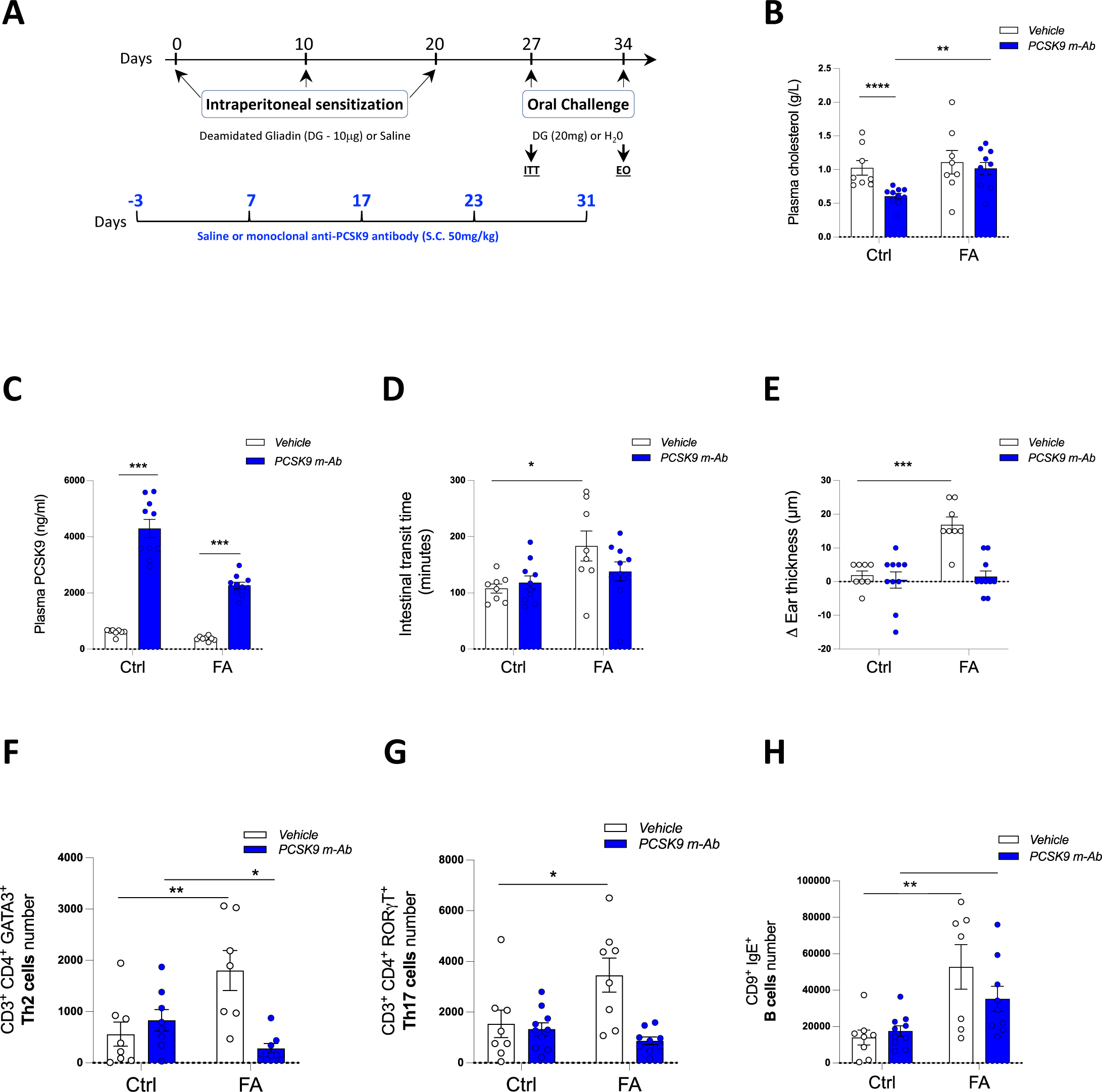
Pharmacological inhibition of circulating PCSK9 significantly reduces gliadin induced food allergic symptoms. (A) Schematic representation of the food allergy protocol: Four-weeks-old BALB/c mice received every ten days a subcutaneous injection of saline solution or monoclonal anti-PCSK9 antibody (alirocumab, 50 mg/kg). Four days after antibodies/saline injection, mice were either sensitized every ten days with intraperitoneal injection of gliadin (DG: 50 µg in 200 µL (PBS + alum); FA group) or saline solution (Ctrl group). Elicitation was induced at days 27 & 34 by oral gavage of gliadin (20 mg in 200 µL of water; FA group). Control mice received an oral gavage of water. Plasma (B) cholesterol and (C) PCSK9 levels were measured at day 34. (D) Intestinal transit time was measured at days 27 following the first elicitation and (E) the ear thickness at day 34 after the second challenge. Th2 (F) and Th17 lymphocytes (G) and IgE^+^ B cells (H) numbers from mesenteric lymph nodes were quantified by flow cytometry. Data are means ± SEM and statistical significance was determined using non-parametric Mann-Whitney *U* test; ns indicates not significant, \**P* < 0.05, \*\**P* < 0.01, \*\*\**P* < 0.001 and \*\*\*\**P* < 0.0001. n=8-10 mice per group.

Taken together, these data indicate that pharmacological inhibition of circulating PCSK9 mimics the same anti-allergic phenotype than those observed in full PCSK9 deficient mice.

### Inhibition of PCSK9 controls food allergy by attenuating the elicitation process

To further determine the phase of the FA process in which PCSK9 inhibition efficiently attenuates the allergic response, we administered PCSK9 m-Ab either only during the sensitization phase (day-3, 7 and 17) or only during the elicitation phase (day 23 and 31) or throughout the entire protocol (**Fig.3A**).

**Figure 3.**
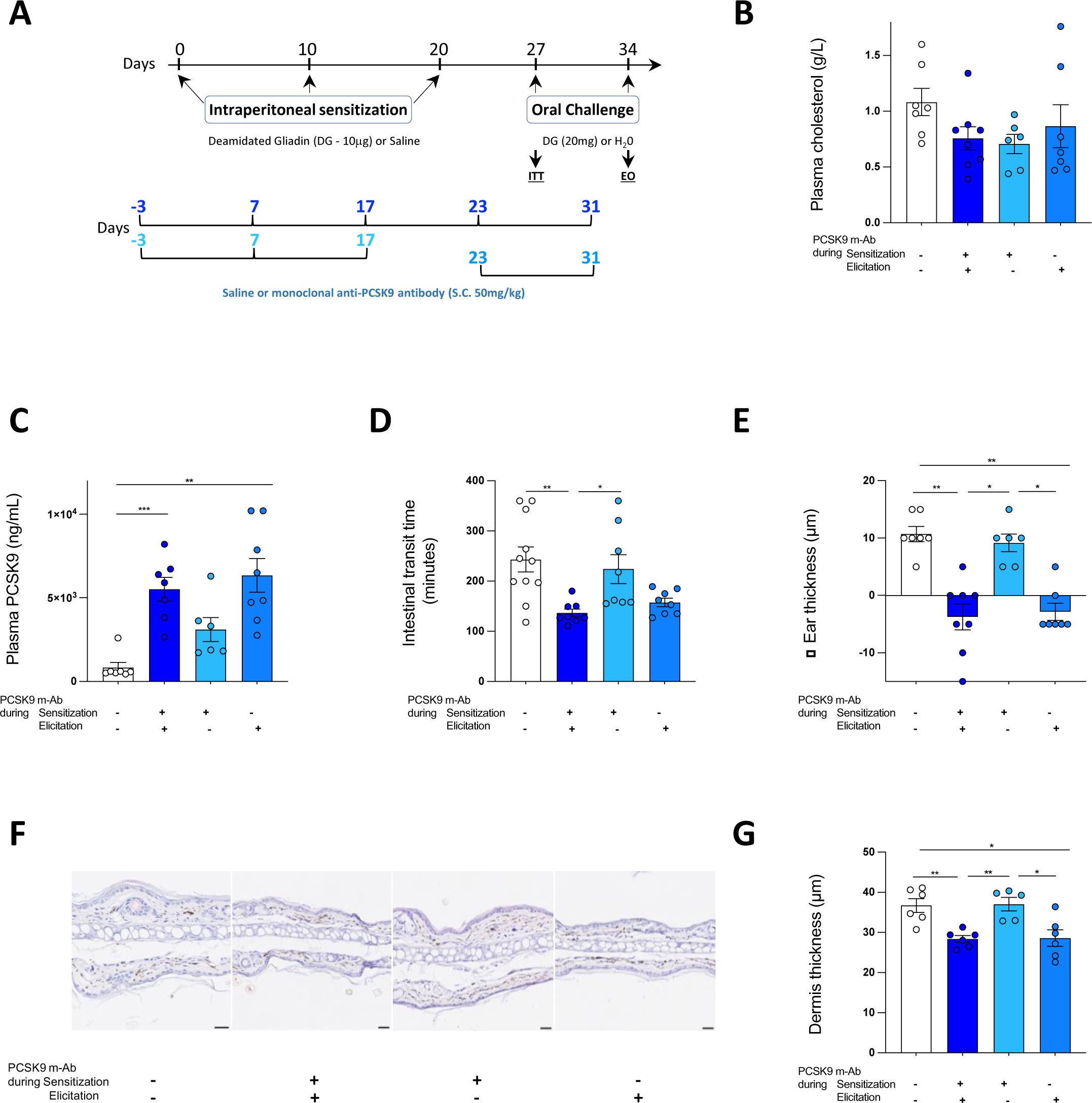
Pharmacological inhibition of PCSK9 during elicitation is sufficient to inhibit food allergy. (A) Schematic representation of the experiment: PCSK9 m-Ab was administrated either during the sensitization steps, the elicitation steps or all along the allergic protocol. (B) Plasma cholesterol and (C) PCSK9 levels measured at day 34. (D) Intestinal transit time and (E) ear thickness measured respectively after the first and second challenge. (F) Representative hematoxylin-eosin-stained ear section harvested 1 hour after the second challenge. Bars, 10 µm. (F) Dermis thickness measurement. Data are means ± SEM and were analyzed using one-way ANOVA; \**P* < 0.05, \*\**P* < 0.01 and *** *P* < 0.001. n=8-11 mice per group.

PCSK9 inhibition during sensitization steps did not prevent transit time induction (**Fig.3B**) or ear oedema (**Fig.3C-E**). By contrast, mice treated with PCSK9 m-Ab during the elicitation phase reproduced the protective effect of PCSK9 inhibition during the entire experimental period (**Fig.3B-E**).

Together, these data suggest that pharmacological inhibition of PCSK9 mostly impacts the elicitation phase and the associated effector cells.

### PCSK9 acts on allergy elicitation process through a LDLR independent pathway

The canonical pathway of PCSK9 action involves the downregulation of the LDLR expression at the cell surface. We aimed to establish whether PCSK9 regulates allergic process induced by FA through a LDLR-dependent pathway. We compared the effect of vehicle and PCSK9 m-Ab treatment on a full *Ldlr* deficiency (*Ldlr^-/-^* mice) and control *Ldlr^+/+^* littermates (**Fig.4A**). The *Ldlr^-/-^* mice present an alteration of plasma cholesterol clearance and cholesterol synthesis control leading to hypercholesterolemia^34^. We did confirm that *Ldlr^-/-^* mice displayed hypercholesterolemia and that PCSK9 m-Ab were unable to reduce their hypercholesterolemia as previously reported^28^ (**Fig.4B**). As shown in **Fig.4C-G**, PCSK9 inhibition efficiently prevented the development of allergic symptoms in both *Ldlr^+/+^*and *Ldlr^-/-^* mice. These results demonstrate that PCSK9 inhibition tunes down DG-FA in an LDLR-independent manner.

**Figure 4.**
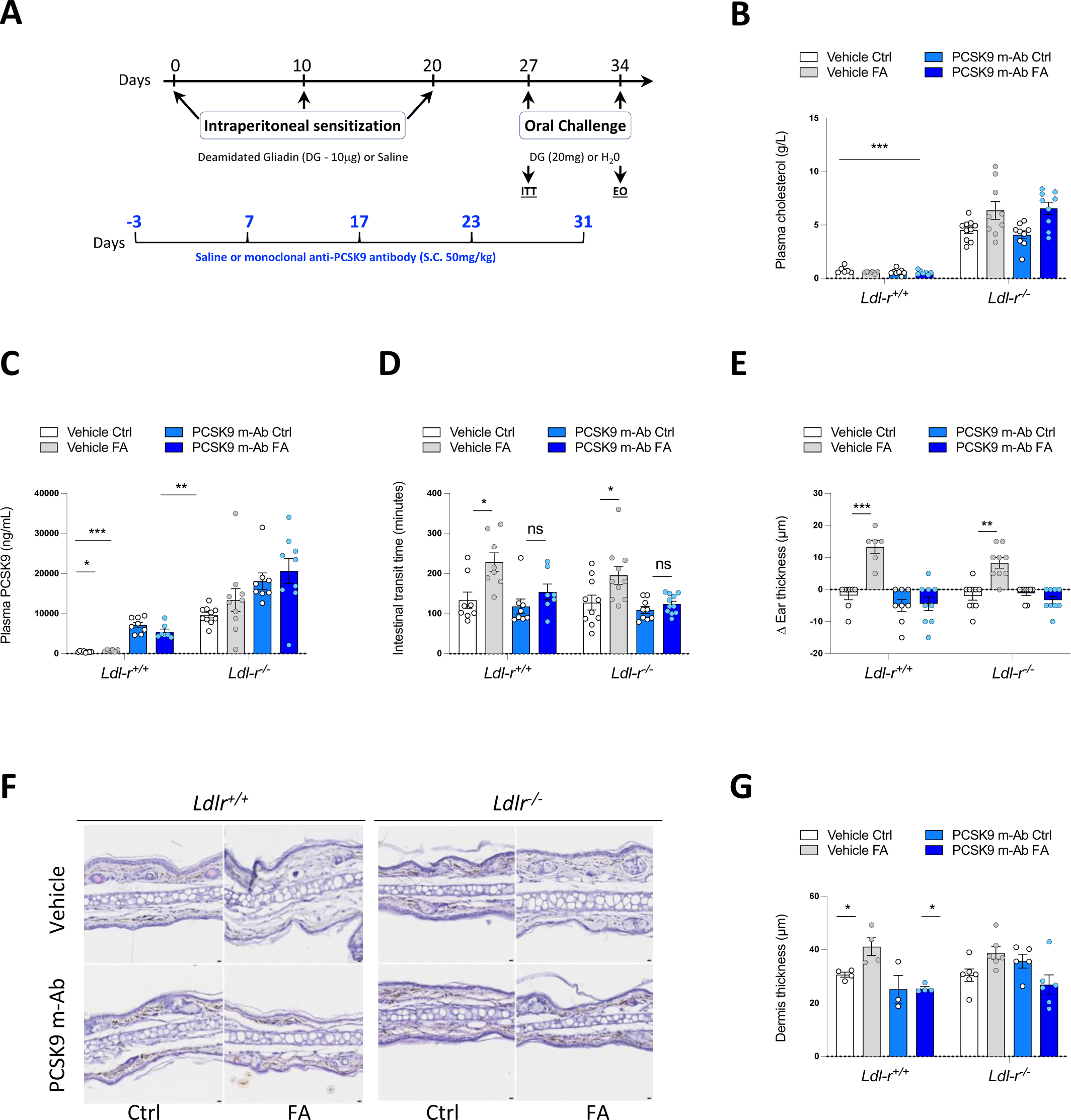
PCSK9 inhibition prevents allergy development in *Ldlr^-/-^* mice. (A) Schematic representation of the experiment: Ctrl and FA *Ldlr^+/+^* and *Ldlr^-/-^* 4-weeks-old *Ldlr^+/+^* and *Ldlr^-/-^* mice were treated with saline solution or PCSK9 m-Ab at day -7, 3, 13, 23 and 33. B) Intestinal transit time and (C) ear thickness measured respectively after the first and second challenge. (D) Plasma cholesterol and (E) PCSK9 levels measured at day 34. (F) Representative hematoxylin-eosin-stained ear section harvested 1 hour after the second challenge. Data are means ± SEM (n= 6 - 10 animals per group). P-values are determined using Mann-Whitney *U* test; ns indicated not significant, \**P* < 0.05, \*\**P* < 0.01 and *** *P* < 0.001.

### PCSK9 affects dendritic cells maturation, CD4^+^ cells viability and basophils degranulation during food allergy

To further characterize the mechanism of action of PCSK9 on FA-mediated immune reaction, we measured the effect of PCSK9 on different target cells types involved in allergic reactions. First, we tested the effect of PCSK9 deficiency on dendritic cell maturation (**Fig.5A**). We showed that *Pcsk9* deficiency does not significantly affect the population of immature or mature dendritic cells (**Fig.5B**). We next tested the functional impact of recombinant PCSK9 treatment on dendritic cells. We did notice that PCSK9 D374Y supplementation in culture media strongly increases mature DC population (**Fig.5C**). Thus, circulating but not intracellular PCSK9 impacts dendritic cells maturation.

**Figure 5.**
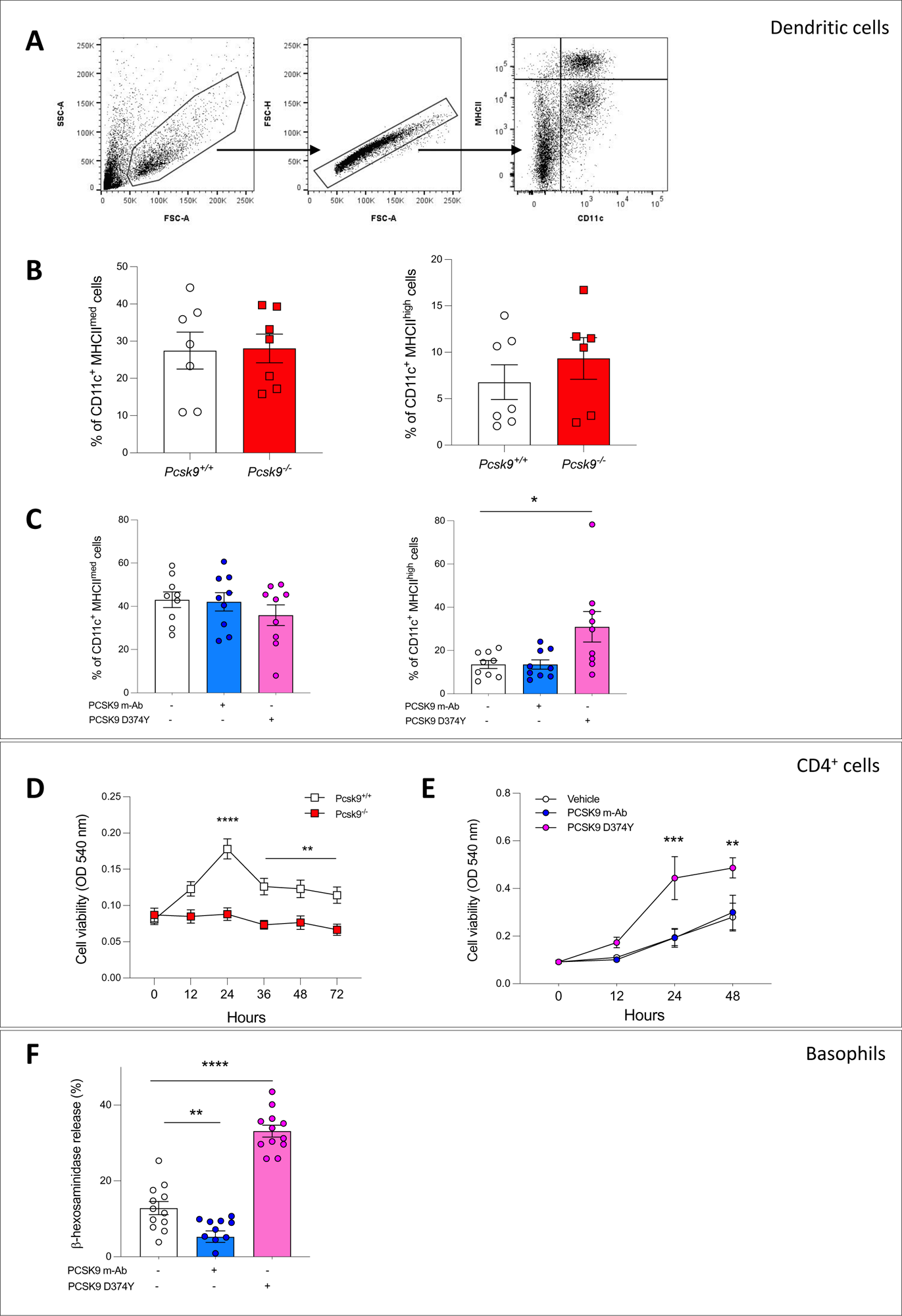
PCSK9 affects immune cells *in vitro*. (A) Schematic representation of the gating strategy (B) CD11c^+^ MHCII^med^ and CD11c^+^ MHCII^high^ cells percentage determined by flow cytometry from *Pcsk9^+/+^* and *Pcsk9^-/-^* BMDCs. (C) BMDCs isolated from *Pcsk9^+/+^* mice were cultured with recombinant D374Y PCSK9 or PCSK9 m-Ab for 8 days, CD11c^+^ MHCII^med^ and CD11c^+^ MHCII^high^ cells percentage was determined by flow cytometry on day 8. (D) Viability of *Pcsk9^+/+^* and *Pcsk9^-/-^* CD4^+^ T cells cultured for 72h. (E) Viability of *Pcsk9^+/+^*CD4^+^ T cells cultured in presence with alirocumab or recombinant PCSK9 D374Y. (F) Basophil (RBL) degranulation was tested by quantifying the β-hexosaminidase percentage release following vehicle, PCSK9 m-Ab or recombinant D374Y PCSK9 treatment. Data are means ± SEM. Mann-Whitney *U* test and two-way ANOVA tests were used; \**P* < 0.05, \*\**P* < 0.01, \*\*\**P* < 0.001 and \*\*\*\**P* < 0.0001. n=5-12 mice per group.

In a second step, we measured spleen CD4^+^ T cell viability from *Pcsk9^+/+^* and *Pcsk9^-/-^* mice. We did observe a lower viability in CD4^+^ cells from *Pcsk9^-/-^*mice compared to those from *Pcsk9^+/+^* mice (**Fig.5D**). Conversely, treatment with recombinant PCSK9 D374Y significantly increased cell viability of CD4^+^ cells (**Fig.5E**).

Finally, we tested the effect of PCSK9 on basophil degranulation. Rat Basophilic Leukemia (RBL) cells were treated with PCSK9 D374Y or PCSK9 m-Ab one hour before gliadin challenge. While PCSK9 m-Ab supplementation significantly reduced β-hexosaminidase release, the addition of recombinant PCSK9 D374Y in culture media strongly promoted gliadin-induced degranulation (**Fig.5F**).

Taking together, our experiments *in vitro* showed that PCSK9 modulates allergy by increasing dendritic cells maturation, CD4^+^ T cell viability and basophil degranulation.

## Discussion

FA is a chronic inflammatory disease due to the persistence of type 2 inflammation at mucosal sites in response to an allergen. The present study is part of the drive to better identify the extrahepatic actions of PCSK9 and in particular its role in immunity. We explored for the first time the action of PCSK9 in FA process and symptoms.

Here, we used deamidated wheat gliadin as food allergen in a well characterized model reproducing the human pathology both by eliciting similar immune reactions^29^ and epitopes^35^. Wheat is a major component of the human diet, particularly in Western countries. Among food allergies, wheat allergy is predominant with an estimated prevalence of 0.8% and 0.1% based on oral challenge experiments in American adults and European populations respectively^2,36^.

We first demonstrated that PCSK9 full deficiency is protective against gliadin-induced FA symptoms occurrence. PCSK9 is predominantly expressed in the liver, kidney and intestine^33^. As food anaphylaxis is mainly characterized by gut symptoms, we explored about the functional role of the intestinal intracellular PCSK9 in immediate allergic reactions. We took advantage of a mouse model generated by our team^28^ which presents a tissue-specific deletion of PCSK9 in the intestinal mucosa. Our data show that these i*-Pcsk9^-/-^*mice remain hypersensitive to allergic symptoms, ruling out the possibility of a major functional role for the intestinal form of PCSK9. Using parabiosis experiments in mice, Lagace *et al.* demonstrated that PCSK9 is secreted into the bloodstream and can act at a distance from its site of production/secretion^38^. We next attempted to decipher the importance of this circulating PCSK9 in FA through its pharmacological inhibition^39^. Our findings clearly indicate that inhibition of circulating PCSK9 confers the same protection as total PCSK9 deficiency.

Although other targets have been identified^40^, the majority of PCSK9 effects on cholesterol metabolism are mediated through its action on the LDLR pathway. In our study, we have nevertheless observed that circulating PCSK9 inhibition in full *Ldl-r^-/-^* mice maintained the same protection as that observed in wild-type mice. If PCSK9 affects certain immune responses via an LDLR-dependent pathway^22,19^, the reverse has also been previously described in the literature. Katsuki S. *et al.* reported that the development of vein graft lesions is facilitated by PCSK9 through mechanisms unrelated to the degradation of LDLR or blood cholesterol levels^32^. More precisely, circulating form of PCSK9 in *Ldl-r^-/-^*mice led to pro-inflammatory macrophage activation leading to immune responses, proliferation and migration^32^. While the mechanisms of action remain relatively unclear, transcriptomic analysis identified NF-κB, lectin-like oxidized LDL receptor 1 (LOX-1) and syndecan-4 (SDC4) as potential pathways of PCSK9 pro-inflammatory role^32^. Along the same line, hyperlipidemic mice due to a double knockout in LDLR and the editing enzyme APOBEC1 feature an increase in Th17 cells and IL-17 secretion in an PCSK9-dependent manner^17^.

How PCSK9 deficiency or inhibition affects FA remains unclear and needs to be further explored. Our *in vivo* data show that several key elements of the allergic reaction cascade are modified by the absence/inhibition of PCSK9. The strong rise in Th2, Th17 and IgE^+^ B cells induced by FA was not observed in *Pcsk9*^-/-^ and PCSK9 m-Ab treated mice. We did confirm these findings *in vitro* by showing that recombinant PCSK9 D374Y 1) promotes the maturation of DC, which may facilitate the initiation of the allergic reaction by modifying the program of naive T lymphocytes in our model; 2) increases CD4^+^ cells proliferation and 3) strongly induces basophils degranulation.

Given the role of PCSK9 in regulating cholesterol metabolism, a relevant question might be whether PCSK9 absence/inhibition exerts an anti-allergic action via modulation of plasma cholesterol concentrations. Our data do not seem to support this hypothesis. We did observe that the pharmacological inhibition of PCSK9 reduces FA symptoms in normocholesterolemic BALB/c mice (**Fig. 3**) but also in hypercholesterolemic *Ldl-r^-/-^* mice (**Fig. 4**). Nevertheless, further data will be needed to determine whether plasma PCSK9 concentrations affect FA by modulating the intracellular and/or membrane cholesterol content of immune cells. Another intriguing point to be explored concerns the loss of regulation of plasma cholesterol concentration in response to PCSK9 in the context of allergy. While the absence/inhibition of PCSK9 was associated with a significant reduction in cholesterol levels under standard conditions (**Fig. 1B & Fig. 2B**), this was no longer the case in allergic situations.

Interestingly, our study shows that PCSK9 inhibition is mostly active during the elicitation phase. More data are required to precisely affirm at which step PCSK9 absence/inhibition exerts an antiallergic effect but it is tempting to speculate that its action on basophils/mast cells degranulation is probably a crucial point. This finding raises the question of the therapeutic value of using PCSK9 inhibitors in allergic crises and to our knowledge, there are no such dedicated studies in the literature. Similarly, there are currently no data showing a potential link between PCSK9 genetic variants and allergy susceptibility.

Thus, our research prospects will focus on understanding the mechanisms of action of PCSK9 on mast cell degranulation induced by wheat gliadin. If we have clearly ruled out the involvement of the LDLR pathway, future studies will be needed to determine which receptors/transporters/membrane proteins can contribute to the attenuation of allergic reaction by PCSK9 absence/inhibition. Indeed, beyond LDLR, PCSK9 has been described as able to interact with other membrane proteins such as the very low-density lipoprotein (VLDLR)^40^, the apolipoprotein E receptor 2 (ApoER2)^40^, the LDLR related protein 1 (LRP1)^40^, CD36^40^, CD81^40^, the β-site amyloid precursor protein cleaving enzyme 1 (BACE1)^40^, the amyloid-like protein 2 (APLP2)^40^, the amyloid precursor protein (APP)^40^, the Na^+^ channel (ENaC)^40^, and (SCD4)^32, 36^. Recently, Liu X. and colleagues found that PCSK9 inhibition can boost the response of tumors to immune checkpoint therapy, through an original mechanism^27^. Indeed, PCSK9 inhibition increases the expression of major histocompatibility protein class I (MHC I) proteins on the surface of tumor cells, promoting a robust intra-tumoral infiltration of cytotoxic T cells^27^. Several studies also highlighted the effect of PCSK9 on the Toll-like receptor 4 (TLR4)/NF-κB pathway ^26,41^. The expression of TLR4 is increased by PCSK9 through the activation of NF-κB pathway, thus, promoting inflammatory cascade^42,43^. Interestingly, TLR4 was associated with food sensitization in mice^44^.

Potential limitations of our study include the fact that our sensitization and allergic reactions in mice were induced by an artificial process, as no spontaneous model of allergic mice is available. Furthermore, only gliadin-induced FA was tested in the present study. It might be of interest to test the effect of PCSK9 on other mouse models of allergic diseases such ovalbumin-induced asthma or cow milk or peanut FA. Next, we did not compare or attempt to combine the anti-allergic effect of PCSK9 inhibitors with anti-IgE immunotherapy^45^. Finally, most of the mouse models used in the present study were bred on a C57Bl6/J genetic background and it is well admitted that is a poor and resistant model of FA in reason of their pro-Th1 immune profile^46^. Reassuringly, most of our findings, including the antiallergic impact of PCSK9 inhibition, was also confirmed on BALB/c mice.

## Conclusion

Taken together, our results highlight the clinical and immunological importance of circulating PCSK9 in food anaphylaxis. Both PCSK9 full deficiency and pharmacological inhibition of circulating PCSK9 by mAbs are protective against the onset of FA symptoms. Our data show that PCSK9 acts mainly during the elicitation phase via an LDLR-independent mechanism, at least by attenuating degranulation of basophils and mast cells.

## Acknowledgment

We thank members of the UTE-IRS-UN animal facility and acklowledge the core facilities Therassay, Micropicell and Cytocell.

## Sources of finding

This study was supported by funding from Amgen Research Protocol Competitive Grant Program (Deciphering the role of PCSK9 in food allergy and asthma), from the Institut de Recherche en Santé Respiratoire des Pays de la Loire, from the Agence Nationale de la Recherche (ANR-16-RHUS-0007: French national project CHOPIN) and from the Fondation pour la Recherche Médicale (INSTINCTIVE research program: EQU201903007846). VL is a recipient of a scholarship from the foundation GENAVIE. The authors have no competing financial interests.

## Nonstandard abbreviations used

APLP2: amyloid-like protein 2
APP: amyloid precursor protein
ApoER2: apolipoprotein E receptor 2
BACE1: β-site amyloid precursor protein cleaving enzyme 1
BMDC: bone marrow-derived dendritic cell
DC: dendritic cell
DG: deamidated gliadin
EnaC: Na^+^ channel
FA: food allergy
FBS: fetal bovin serum
GM-CSF: granulocyte macrophage colony stimulating factor
Ig: immunoglobulin
IL: interleukin
LDL-c: low-density lipoprotein cholesterol
LDLR: LDL receptor
LOX: oxidized LDL receptor
LRP: LDLR-related protein
MHC I: major histocompatibility protein class I
MLN: mesenteric lymph node
PCSK9: Proprotein convertase subtilisin/kexin type 9
RBL: rate basophilic leukemia
SDC: syndecan
TLR: Toll like receptor
VLDLR: very low-density lipoprotein receptor

## Highlights

- Both pharmacological inhibition and *Pcsk9* deficiency protect mice from food allergy.
- PCSK9 inhibition is mainly effective during the allergic elicitation phase
- PCSK9 acts on allergy through a mechanism independent of LDLR, at least by modulating degranulation of basophils and mast cells.

## Notes

### Competing Interest Statement

The authors have declared no competing interest.

